# Scars of hope and forewarn of a bleak future: Post-mortem findings of dugongs (Dugong dugon) belonging to a relict population in the Gulf of Kachchh, India

**DOI:** 10.1101/2021.12.21.473634

**Authors:** Sameeha Pathan, Anant Pande, J.A. Johnson, Kuppusamy Sivakumar

## Abstract

A relict dugong population resides in the Gulf of Kachchh in Gujarat state of India. Very little is known on this population stemming from lack of focused studies and inadequate examinations of previous strandings. In this study, crucial ecological information is gathered through a systematic necropsy on stranded dugongs in the gulf. As indicated through dorsal tusk-rake scars on both the carcasses, this study presents first records of derivative physical evidence to the presence of a socially interacting population. Progressive healing and differences in width of the scars indicated more than two individuals had participated in a sexual agnostic or courtship event. Conversely, our findings report that both the animals suffered relative effects of asphyxiation after fishing net entanglement. An implication of a prior pathological condition(s) in the form of dermal cysts, swollen mesenteric lymph nodes, and endoparasites are also reported. Stomach content was examined for a qualitative dietary characterization. Other potential threats along with fishing net microfilaments found in the stomach contents of both dugongs are discussed in brief.

## Introduction

Indian dugong populations are considered to be “Regionally Endangered” as compared to the species’ global IUCN red list status of vulnerable^1^ making them the most threatened marine mammal species along the Indian coastline. Their foraging range along the shallow near-shore areas brings them in direct conflict for space use with artisanal fisheries and other coastal activities such as tourism, port activities, etc. The current range of dugongs in the country is restricted to parts of the Gulf of Kachchh (Gujarat), Palk Bay, Gulf of Mannar (Tamil Nadu), and Andaman & Nicobar Islands (across the islands except for Great Nicobar). Several research advances related to the Tamil Nadu and Andaman and Nicobar Islands’ dugong populations pertaining their foraging behaviour ^2^, abundance^3,4^ etc. There is a significant lack about the ecological background information regarding dugong population of the Gulf of Kachchh (GoK).

To date, no extant dugong population has been confirmed from the west coast of India except for the small population of the GoK, making it an important dugong habitat in the west coast of India. This population is known to be restricted to the islands and reefs of the southern coast of the Gulf ^5^. A natural low fecundity and slow growth rate make dugong populations around the world extremely sensitive to adult mortalities^6^. Further, the low population size and in general elusive nature of dugongs make them difficult to study in the wild. For a small marine mammal population, studying the type and consequences of anthropogenic pressures becomes a difficult feat in wild. Given the lack of long-term datasets on this geographically isolated population, stranding events provides a rare window to collect essential data about health, feeding ecology, and reproductive status etc which otherwise remains undetected due to the cryptic nature of some animals. The extent of information salvaged from dead strandings depends upon the stage of carcass decomposition. We present a brief report on salvaged information deduced from two dugong carcasses of stage I and stage IV decomposition state

## Methods

We obtained dugong stranding information through a volunteer network developed in the fisher community of Beyt-Dwarka, and Arambada village of the GoK. This network comprised of fisherfolk who were sensitized through a series of community interaction programs (n = 3) conducted at Okha and Beyt-Dwarka fisher villages; Balapur, Arambada, and Rupen. Two dugong strandings were reported in the months of February 2018 and May 2018. Initial picture of Ajad island dugong was taken when the carcass was relatively fresh (Fig1.1). The carcass was reported, and was necropsied after two weeks of the stranding event. Both necropsy examinations were done on-site under the supervision of the Gujarat state forest department.

Carcass photo-documentation and necropsy were performed using standard salvage and necropsy procedures^7^. Each animal was examined for external marks, lesions, bruises and other injuries prior to internal examination. Since decomposition is relatively faster in intestines, Ajad dugongs’ visceral organs after stomach, were rendered unexaminable.

Stomach content analysis was conducted on samples (300 g) collected from the cardiac end. The stomach content was preserved using 10% neutral-buffer formalin within nine hours of necropsy. Qualitative analysis of the gut content was done after diluting 5 grams of sample (n=2) with 50 ml of distilled water. To avoid crowding of seagrass segments, from the diluted sub-sample (50ml), a volume of 5 ml was used for making observations on a 2×2 cm^2^ graded petri-plate.

## Results

### Ajad island stranding

A stage four ‘badly decomposed’^7^ carcass was necropsied at the north-eastern part of Ajad island (center point-22°23’ N, 69°19’ E), where it was stranded (Fig1.2). The animal was a 2.6 m (straight body length) adult female. The skin had been sloughed off completely but the remaining attached skin on the mid-dorsal side retained a hyperpigmented rake mark (Fig1.1), evident of the fact that it had interacted with other adult male dugongs. The head showed clear signs of severe intramuscular haemorrhage with a cutaneous abscess on and around the nostrils (Fig1.1). This haemorrhage could result from blunt force trauma to the head after a boat collision. On the dorsal side of the animal, the caudal vertebral processes were markedly visible (black arrow, Fig 1.1) indicating emaciation, but other signs of starvation such as caved in body mass around the peduncle could not be verified due to advanced stage of decomposition. Starvation was further ruled out as the stomach was full of seagrasses and blubber layer (Fig1.5) was intact in both consistency and thickness. A cross-section examination of the dermal wall of the thorax (Fig 1.3) revealed the presence of a single oval and well-demarcated parasitic cyst of (dimensions; 32.11 × 40.40 mm) (Fig 1 (4)). Nematode, *Paradujardinia halicoris*, a common endoparasite in dugongs ^8,9^, and manatees^10^, was found in the stomach of the animal.

**Figure 1.**
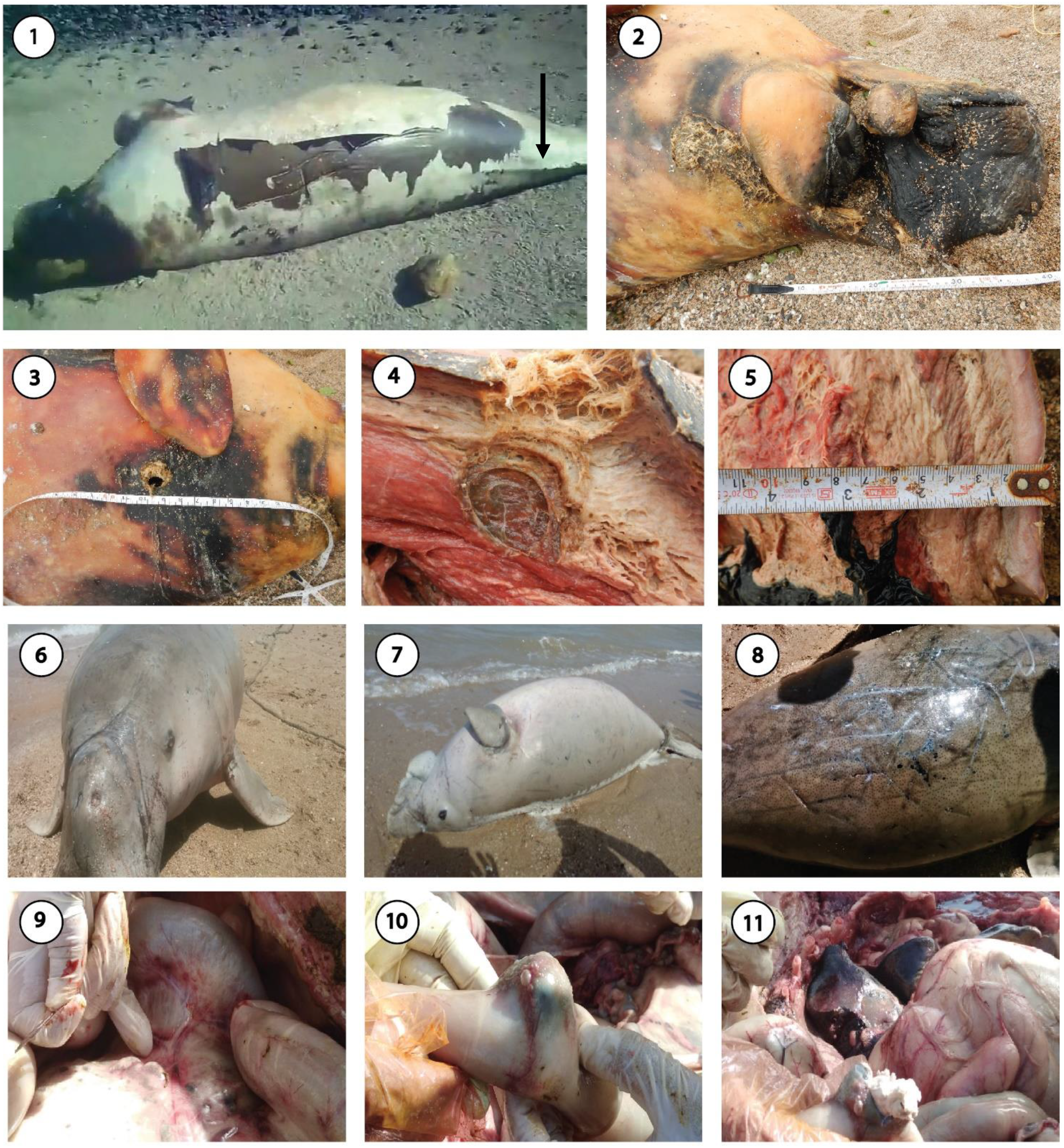
Ajad dugong stranding; (1) Decomposed carcass of a mature female dugong (2) Rope burn marks around the neck (3) thorax with cavity caused by parasitic infestation (4) Cross section through the cavity showing cyst of an unidentified parasite (5) Cross section across the mid-ventral tissue layer Positra dugong stranding; (1) Positra dugong carcass (2) Rope burn marks visible around the snout (3) Numerous healed wounds on the back (4) swollen mesenteric lymph nodes (5) & (6) Purulent tumour in the small intestine

### Positra island stranding

A sub-adult male dugong of the straight-body length of 2 m was reported by fishers of Positra near Man-marudi Island (center point-22°25’59.56”N, 69°13’23.86”E) of the GoK. There were no signs of decomposition (bloating, discoloration, etc). Since the flippers had mobility when the carcass was found, rigor mortis had not set in, conclusively indicating mortality in the past few hours. Fishers confirmed mortality due to suffocation after net-entanglement which was factualised by rope-burn marks around the head and neck area (Fig 1.6). The skin was in a good condition and showed no signs of sloughing (Fig1.7). The intact skin indicated various important clues to the type of interaction the dugong had undergone before its death. Small scars of several healed wounds (Fig 1.8) were found extending from the lower neck to the peduncle of the dorsum of the animal, the significance of which is discussed later.

Since the visceral organs were intact, a thorough examination of the gastrointestinal tract could be conducted. A localized single tumour on the duodenum which was sized 5 cm radius was found (Fig1.9). The outgrowth’s purulence had a hard greasy consistency (Fig1.10). Moreover, the mesenteric lymph nodes appeared swollen (Fig1.11).

### Tusk-rake scars

Several healed indentations and wounds were found on the dorsum of the animal from various interactions (Figure 2). The source of these wounds can be either biological via intraspecific interactions or non-biological source made by scratching behaviour of dugongs over sandy sediment^11^. Except for a few deep indentations, most of these marks were shallow. More important are the parallel marks on the back which are made due to the grazing action of pair of tusks. We discovered several counts of such *tusk-rake scars* (TRS) on the back of this dugong individual. Scars with different widths indicated that sexually mature conspecifics of different age-group had interacted with this male dugong (Fig 2.1). Based on width, five distinctly different scars were identified. These were then broadly categorized into two stages based on the progressiveness of healing (Fig 2.2). A new scar indicates a ‘recent’ (Stage -1) agnostic interaction, whereas completely healed marks appear shallow and discolored^12^ are evidence of ‘older’ (Stage-2) instances. Out of these, only one was recent looking (Stage 1) as it hadn’t completely healed (scar 1 in Fig 2.1). The rest of the four scars (scars 2-5 in Fig 2.1) were older (Stage-2) and are probable to be made during a single event. Hence, at least two separate events of ritualistic sexual conflicts were experienced by the Positra dugong. Moreover, a scar inflicted by a relatively younger dugong (mark 4, width-5.3 cms) was also observed.

**Figure 2.**
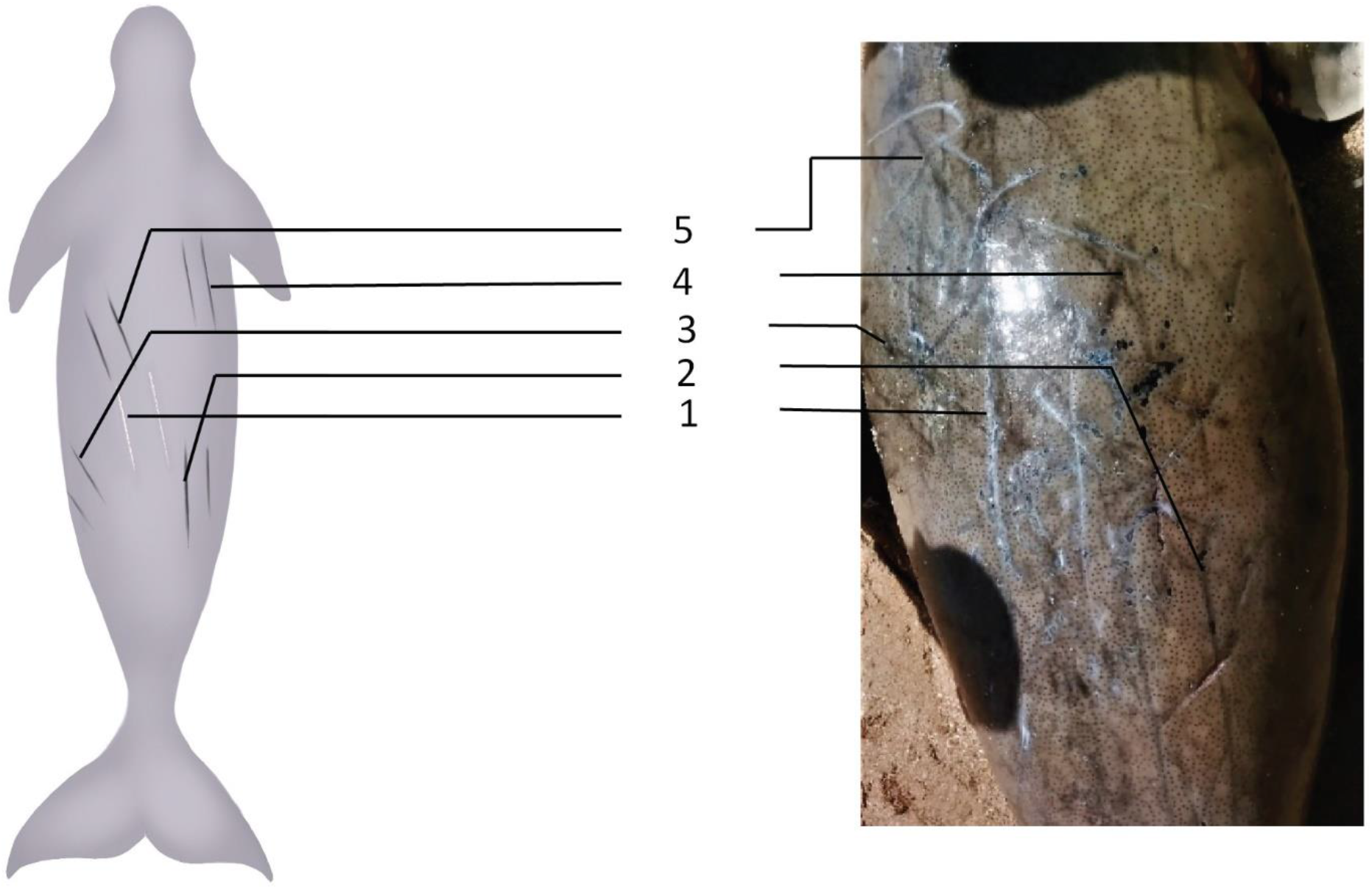
An illustration of tusk rake marks (TRS) as seen on the dorsum of dugong in Positra

**Fig 3.**
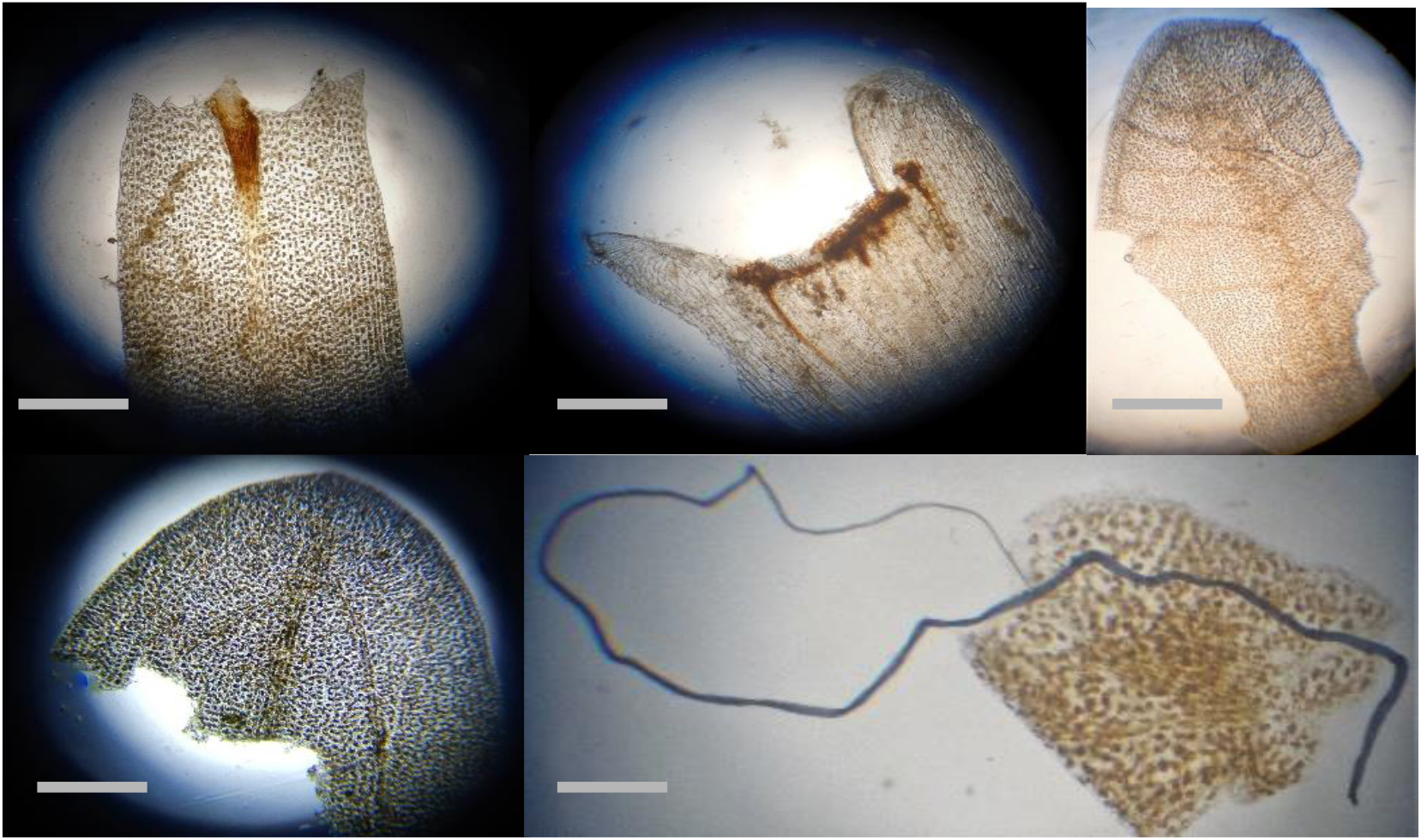
Microphotographs of seagrass and plastic micro-filaments from the stomach content (scale-2mm). (1) *Halodule uninervis* leaf apex (2) *Halodule uninervis* leaf sheath (3) *Halophila beccarii* (4) *Halophila ovalis* (5) Nylon micro-filament from fishing nets

**Table 1.** Measurements and healing stages of six different tusk rake marks observed on dugongs stranded in the Gulf of Kachchh, Gujarat, India. TRS6’s healing stage i.e., recent or a healed scar was not clear from the photographic evidences

| Tusk rake scars (TRS) | Dugong case | Length (in cms) | Width (in cms) | Healing stage |
| --- | --- | --- | --- | --- |
| TRS1 | Beyt- Dwarka | 33.3 | 11.3 | Stage 1 |
| TRS2 | Beyt- Dwarka | 26.5 | 8.7 | Stage 2 |
| TRS3 | Beyt- Dwarka | 22 | 6.5 | Stage 2 |
| TRS4 | Beyt- Dwarka | 18 | 5.3 | Stage 2 |
| TRS5 | Beyt- Dwarka | 21.4 | 10 | Stage 2 |
| TRS6 | Ajad | 32 | 10.8 | NA |

**Table 2.** Body measurements of the two dugong individuals stranded in Gulf of Kachchh, Gujarat, India.

| **Measured part** | **Ajad dugong (in cms)** | **Positra dugong**<br>**(in cms)** |
| --- | --- | --- |
| Straight body length | 260 | 200 |
| Girth (from under flippers) | - | 138 |
| Width (from and to the base of flippers) | 85 | 61 |
| Right flipper (dorsal proximal end to the tip of the flipper) | 42.5 | 15 |
| Left flipper | 36 | 14.7 |
| Maxilla | 20 | 16.5 |
| Teat length | 6.5 | - |
| Fluke length | 88.7 | 78 |

### Stomach content analysis

Leaf tip characteristics, leaf venation patterns, and numbers, leaf scars on rhizomes are the most prominent and important morphological features examined to identify seagrass species^13^ (Fig 5); *Halophila beccarii, Halophila ovalis*, and *Halodule uninervis* were found in the stomach contents of both the stranded individuals. Both dugongs had also ingested fragments of nylon fishing net micro-filaments of length range; 0.23–30 mm

## Discussion

There is very little information regarding the social and reproductive behavior of dugongs in India. As concluded from the presence of tusk rake marks, both dugongs were involved in inter and intra-sexually aggressive conflicts which reflect positively on the social behavior of this small population in the Gulf. Similar teeth rake marks in bottlenose dolphins are reported to be a very reliable indicator of conflicts over resources and mates^14^. Paired parallel in dugongs also provided insights to the diversity in age-class of the participant dugongs in the Gulf of Kachchh.

Dugong cows that are less than 2 mts long have almost certainly not born a calf (non-parous), whereas those larger than 2.50 m are likely to have been born young (parous)^15,16,17^. Hence, conclusively, the Ajad dugong was a sexually mature female, although due to the liquefaction of the ovaries, it could not be confirmed whether the cow had birthed a calf during her lifetime or not. Marine traffic exerts a growing pressure on marine megafauna^18^. This contributes to a net-entanglement as one of the leading causes of death for dugongs^19,20^. Dugong population in the GoK, although range-resident, are highly mobile within the range. Their movement response is adapted to a dynamic tidal regime and local knowledge of the widely distributed seagrass habitats in the Gulf. This makes them mildly susceptible to harm caused by fishing activities that coincide with their foraging habitats.

Dugongs, like other marine mammals, are susceptible to a wide range of diseases, some of the infectious, non-infectious, or idiopathic^7^. Both stranded animals, collectively, suffered from underlying conditions like visceral tumors, inflamed mesenteric lymph nodes, and parasitic cysts. This evidence might indicate the body’s immunoreactive state before its ultimate death. Our study reports that although the Ajad dugong might have suffered from head trauma due to boat collision and underlying parasitic infection. This can further be consolidated as the animal showed distinct signs of emaciation, yet had a relatively full stomach. A fishing net entanglement is evident from the rope impressions around the neck and is suspected to be the ultimate cause of its death. The same can be concluded for the Positra dugong.

Seagrass meadows are known to reduce the velocity of tidal currents^21^, thereby causing sedimentation of fine particles^22^ and co-incidentally of fishing net micro-filaments. This naturally makes seagrass meadows extremely prone microplastic pollution, herbivory of which becomes a medium for plastic/nylon micro-filaments to enter to diverse food chains. Anthropogenic marine debris has also been documented in subtidal seagrass meadows of the Philippines^23^. However, the impact of micro-plastic sedimentation in dugong foraging habitats depends upon their bio-availability and post-consumption consequences in dugongs, which remains obscure. Although the clinical significance of ingestion of plastic micro-filaments is unknown in dugongs, it can have potentially lethal effects on sediment biota of seagrasses.

Pigeye shark, Pelagic thresher shark, Black tip reef shark, etc are amongst the large sharks that inhabit the GoK waters seasonally^19,23^. Whether dugongs are at predation risk by these migratory sharks remains uncertain as to date no dugong carcass retrieved have reported any shark bite marks in the Gulf. However, anecdotal data acquired from interviews with fishermen suggest targeted shark fishing during the monsoon season might be a potential pressure on the survival of dugongs in the GoK. During the monsoon fishing ban period, sharks are occasionally and illegally sought after and are considered a valued catch. It has been suggested that the meat of dugongs, dolphins, and batoids are sometimes used as bait to lure commercially important sharks.

Photo-documentation and necropsy techniques as one of the key conservation tools to understand underlying stressors to the cryptic dugong population. Studying foraging behavior and reproductive status of dugongs in wild is close to impossible, especially for a small population wherein the chances of detecting a live animal become rare. This obscurity is further complemented by the turbid waters of the Gulf (0.5-2 mts visibility), which reduces the detection probability of the animal. Hence, an idea about their biology, reproductive status, and health, and natural and anthropogenic stressors can only be understood with relative clarity by salvaging information from carcasses.

Partly, the initiation of our research regarding dugong foraging ecology across the Gulf ^25^ was an extension to conclusions drawn from the necropsy examinations. GoK is home to a variety of species that are ecologically significant and share the same habitat as dugongs. With dugongs being an umbrella species, their conservation may indirectly protect these ecological communities.

## Acknowledgement

We thank the Gujarat Forest Department for the arrangement of necessary provisions needed to conduct and supervise the necropsy procedures and the funding agency; National CAMPA Advisory Council (NCAC), Ministry of Environment, Forests and Climate Change, Government of India. We also thank field assistant Junus Babbar and Sattar Nagiya for their relentless support as key informants and Vabesh Tripura for the illustration and figure presentation.

## Declarations

### Authors’ contributions

Sameeha Pathan – Writing (Original draft preparation), Reviewing, Editing, Conceptualisation, investigation, methodology, and visualisation

Anant Pande- Editing, project administration

J.A. Johnson, K. Sivakumar- Funding Acquisition, project administration, supervision

### Ethics approval

No approval of research ethics committees was required to accomplish the goals of this study as the necropsies were performed under the guidance of veterinarians and officers from the Gujarat Forest Department, Government of India.

